# Expression of endoplasmic reticulum mannosyl-oligosaccharide α1, 2-mannosidase regulates tumorigenicity of prostate cancer cells

**DOI:** 10.1101/750984

**Authors:** Christian Toonstra, Xiangchun Wang, Naseruddin Höti, Jianbo Pan, Hui Zhang

## Abstract

It has been previously demonstrated that glycogenes can function as important pleiotropic regulators of tumorigenesis. In this study, we investigated the function of endoplasmic reticulum mannosyl-oligosaccharide 1, 2-alpha-mannosidase (MAN1B1) on the behavior of prostate cancer cells and found that overexpression of MAN1B1 in androgen-independent PC3 cells reduced cellular migration. We extended the analysis to an androgen dependent cell line, LNCaP, and another androgen independent cell line, LNCaP-AI, observing a similar migratory phenotype. By using quantitative proteomics, we found the downregulation of vimentin expression when PC3 or LNCaP-AI cells with overexpressed MAN1B1. The current study suggests that MAN1B1 may regulate cellular migration and promote epithelial-to-mesenchymal transition (EMT) phenotype in prostate cancer cells.

## Introduction

In spite of intensive research and significant advances in the understanding of prostate cancer (PCa) pathogenesis, PCa remains the most commonly diagnosed cancer in American men, and is one of the major causes of cancer-related mortality (1). Traditional treatment options for organ-confined PCa, including radical prostatectomy and localized radiation therapy are hampered by systematic recurrence in some patients (2). In advanced prostate cancer, the gold-standard treatment, androgen-deprivation therapy, is eventually overcome by PCa progression to the castration resistant phenotype (3). The conversion of PCa to a more aggressive phenotype is a multistep process involving an unknown number of cellular alterations. A clearer understanding of the changes that occur in the progression of localized, non-aggressive PCa to a more aggressive, androgen independent (AI) phenotype will potentially provide additional insight into treatment options by opening cellular pathways that can be targeted for treatment to suppress tumorigenesis.

A number of studies have reported pathways, including cell proliferation (4,5), cell death (6), and alterations of extracellular matrix (ECM) adhesion proteins (7,8), that are critical steps in PCa tumorigenesis and are promoted by the loss or gain of regulatory proteins. The tumorigenesis process relies on a series of sequential, obligatory steps beginning with the detachment from neighboring cells and the concomitant degradation/remodeling of the basement membrane, both of which are barriers to dissemination (9,10). The second step, intravasation, requires the tumor cells to disrupt the cell-cell junctions that seal the lumina of blood vessels (11,12). Both pathways are required for PCa cells to separate from the primary tumor site and enter the circulatory system, migrating to a target organ and forming secondary metastatic colonies. A critical step in PCa progression is the epithelial-mesenchymal transition (EMT), in which epithelial cells acquire an altered phenotype, characterized by enhanced migratory and invasive potential. The conversion of polarized epithelial cells to the mesenchymal cell phenotype, facilitated by EMT, occurs via a series of biochemical alterations that ultimately lead to the release of the cell from the basement membrane (13).

Expression levels of glycosylation-related enzymes are known to be disrupted during tumorigenesis (14, 15). Indeed, aberrant glycosylation in cancer progression can be used to distinguish cancerous cells from healthy cells, and is one of the few distinctive surface details that can be used to distinguish between self-derived antigens (16,17). As a non-template mediated post-translational modification, glycans are not under direct genetic control, however, the pattern of glycosylation is indirectly determined by the expression levels of glycosyltransferase and exo-/endo-glycosidase enzymes, pathways known to be dysregulated by cancer (18, 19). Glycoproteins on the cell surface or secreted to the extracellular space are extensively processed to complex or hybrid-type glycans and oligomannose glycans are less abundant in normal serum and tissues. Previous reports have demonstrated the increase in the abundance of oligomannose glycans in several cancer types (20–26). The presence of increased levels of oligomannose glycans in cancer represents an aberration in the biosynthetic pathway of protein glycosylation in cancer cells. Indeed, oligomannose-reactive antibodies (Abs) have been isolated from patients with late-stage PCa (27), suggesting that oligomannose trimming is dysregulated at a stage within the glycan processing pathway. The observation that the abundance of oligomannose glycans is increased during tumorigenesis suggests that there is dysregulation of the enzymes responsible for trimming mannose during glycan biosynthesis. The pathological dysregulation of oligomannose-trimming enzymes could be related to the enhancement of tumorigenesis.

MAN1B1 encodes an endoplasmic reticulum-associated α1, 2-mannosidase responsible for the specific release of a single mannose monosaccharide from the middle branch (D2 arm) of Man_9_GlcNAc_2_ yielding the Man_8_GlcNAc_2_ regioisomer B (Man8B) (Scheme 1) before Man_8_GlcNAc_2_ migration to the Golgi. MAN1B1 occupies a central position in the glycan biosynthesis pathway as it is the first step in the trimming of oligomannose (high-mannose) glycans (28). Downregulation of MAN1B1 expression leads to decreased glycan processing resulting in a higher abundance of oligomannose glycans and a decrease in the abundance of both complex and hybrid-type glycans. Indeed, a recent study demonstrated that knocking out MAN1B1 alone was enough to reduce the expression of most complex-type glycans in HEK293 cells (29). We speculated that, given its central role in N-glycan processing, overexpression of MAN1B1 could yield a significant alteration in the glycan phenotype on the cell surface. The change in cell-surface glycan phenotype following MAN1B1 overexpression could influence the ability of the cells to adhere to the cellular milieu and potentially minimize metastatic potential.

In the present study we investigated the differential expression of endoplasmic reticulum (ER) mannosyl-oligosaccharide 1, 2-alpha-mannosidase (MAN1B1), the initial mannosidase encountered by nascent proteins within the ER, in PC3 PCa cells. We observed a dramatic reduction in the migratory phenotype following MAN1B1 overexpression. We extended the study to two other PCa cell lines, LNCaP and LNCaP-AI cells, finding that overexpression of MAN1B1 consistently produced a reduction in cell migration across all three cell lines, suggesting that the expression of MAN1B1 may play a critical role in PCa tumorigenesis. To investigate the molecular changes that might be responsible for the MAN1B1 function in prostate cancer cells, we applied a global proteomics approach to the wild-type and MAN1B1 overexpressed cells of all three cell lines and found that MAN1B1 overexpression resulted in the downregulation of vimentin expression in protein levels in androgen-independent PC3 and LNCaP-AI cells, suggesting that MAN1B1 may regulate cellular migration and influence the epithelial-to-mesenchymal transition (EMT).

## Results

### Overexpression of MAN1B1 is associated with a non-aggressive phenotype of prostate cells

MAN1B1 is the first mannosidase encountered by nascent glycoproteins, and alterations in MAN1B1 expression could have an important downstream impact on cellular behavior. Given the observation that oligomannose glycan expression is more abundant in tumor cells compared to normal cells in number of cancer types (20–25, 27, 30), we became interested in examining the function of MAN1B1 in PC3 cells. PC3 cells are a PCa cell line with high metastatic potential (31). Overexpression of MAN1B1 potentially result in a higher abundance of hybrid or complex-type glycans in the transformed cells, which may influence tumor cell behavior. Stably MAN1B1-transfected PC3 cells were generated using cDNA, and overexpression of was confirmed by quantitative global proteomics using isobaric tags for relative and absolute quantitation (iTRAQ) labelling. MAN1B1 protein expression was found to increase 2.7-fold, compared to the WT PC3 cell line, based on proteomic iTRAQ quantitation (Figure 2A).

**Figure 1.**
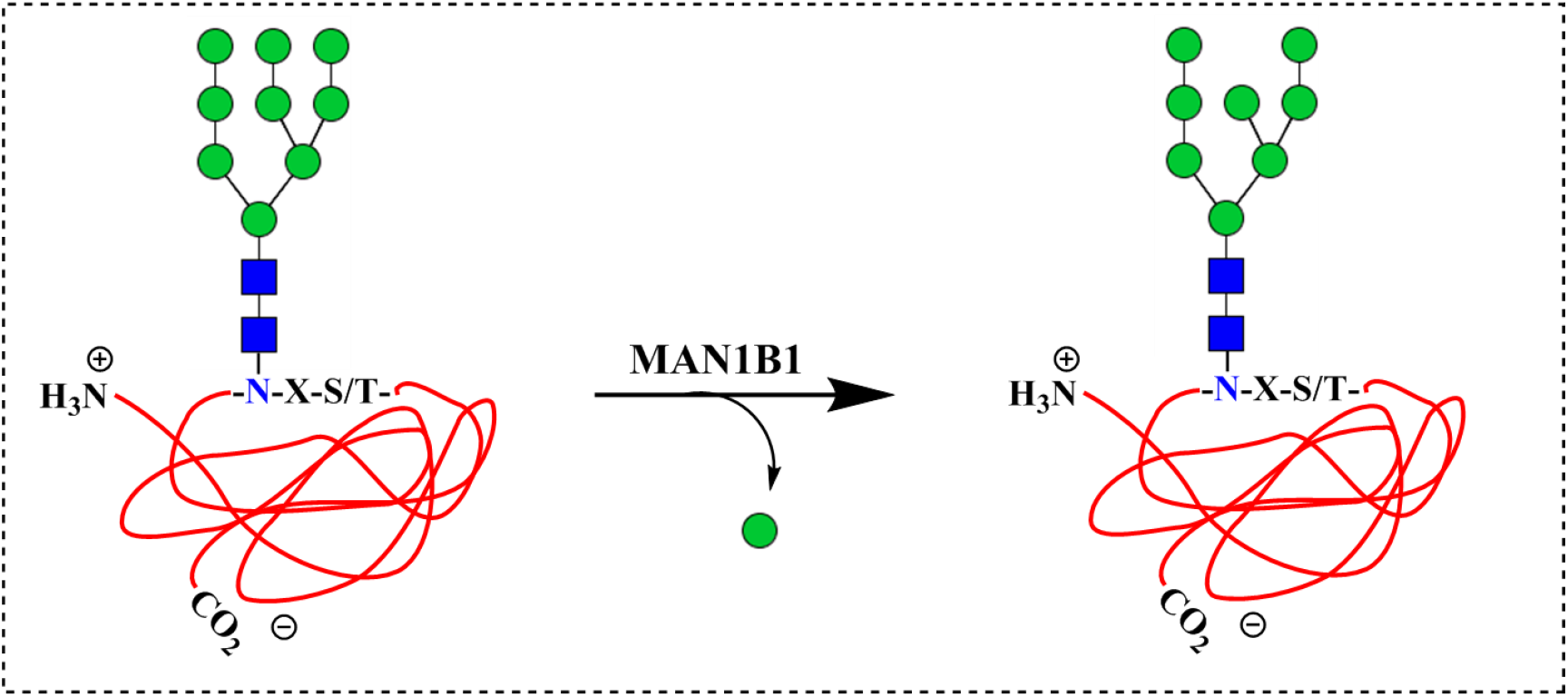
ER α-1,2-mannosidase selectively cleaves the α1, 2-Man-Man linkage from the terminal D2 arm of nascent folded proteins prior to export to the Golgi apparatus.

**Figure 2.**
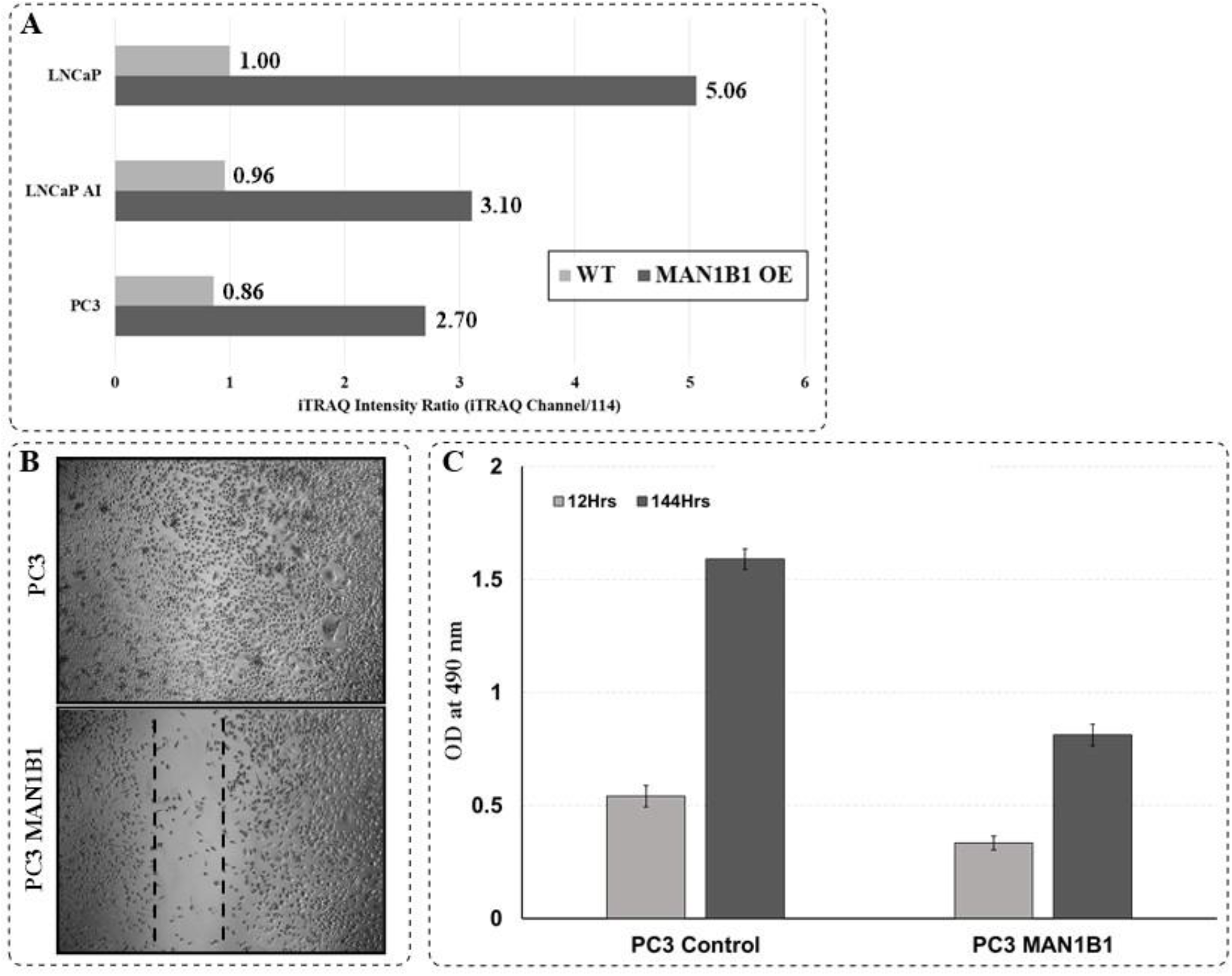
Overexpression of MAN1B1 in prostate cancer cells is associated with a less aggressive phenotype. **A)** Expression level of endoplasmic reticulum mannosyl-oligosaccharide 1, 2-alpha-mannosidase (MAN1B1) in different PCa cell lines with and without MAN1B1 overexpression. **B)** Scratch closure assay comparing PC3 (WT) and PC3-MAN1B1 cell lines. PC3-MAN1B1 exhibited a reduced scratch closure rate compared to PC3 (WT) cells. Zone clearance is 500 μm. **C)** MTS Assay indicated that MAN1B1-PC3 cells demonstrated lower proliferation compared to PC3 (WT) cells.

To understand if MAN1B1 overexpression has biological significance in PCa cellular aggression, we performed a series of scratch closure assays in the confluent monolayer of cultured PC3 cells. Our data indicated that MAN1B1-expressing stable PC3 cells (i.e. PC3-MAN1B1) exhibited a retarded scratch closure rate compared to the control (WT PC3) cells (Figure 2B). The scratch closure assays indicate that increased expression of MAN1B1 reduces cell migration and may contribute to a less aggressive phenotype.

Upon observing the reduction in aggression, indicated by the decrease in PCa cell motility in PC3 cells following MAN1B1 overexpression, we investigated the impact of cell proliferation by MAN1B1 by performing an MTS assay. The viable cell number for both PC3 and PC3-MAN1B1 proliferation increased in a time-dependent manner, however, WT PC3 cells exhibited higher proliferation rates compared to PC3 cells that overexpress MAN1B1 (Figure 2C). The decrease in cell proliferation for PC3-MAN1B1 cells was particularly dramatic with time, with the PC3 cells showing nearly 2-fold higher proliferation rates compared to PC3-MAN1B1 cells after 144 hrs (Figure 2C).

### Impact of MAN1B1 overexpression on the migration behavior of LNCaP and LNCaP-AI cell lines

Following the initial observation that overexpression of MAN1B1 alone in PC3 cells is sufficient to reduce cellular migratory behavior, we chose to analyze two additional PCa cell lines with low to moderate metastatic potential, including the androgen-dependent LNCaP cell line and androgen-independent LNCaP-AI cell line (32, 33). Analysis of additional cell lines will help to establish the migration phenotype as a general result of MAN1B1 expression across several PCa cell lines.

The impact of MAN1B1 overexpression on cellular migration in LNCaP and LNCaP-AI cell lines was determined by scratch closure assays (Figure 3A, B). Although LNCaP (WT) demonstrated slow migration behavior, the LNCaP-AI (WT) cell line exhibited significant migratory capacity (Figure 3B). Following MAN1B1 overexpression, both LNCaP-MAN1B1 and LNCaP-AI-MAN1B1 demonstrated a reduction of migration capacity (Figure 3A, B). The migratory phenotype of LNCaP and LNCaP-AI upon MAN1B1 overexpression was similar to the phenotype observed for PC3 cells, suggesting that MAN1B1 overexpression can reduce cellular migration across three PCa cell lines tested.

**Figure 3.**
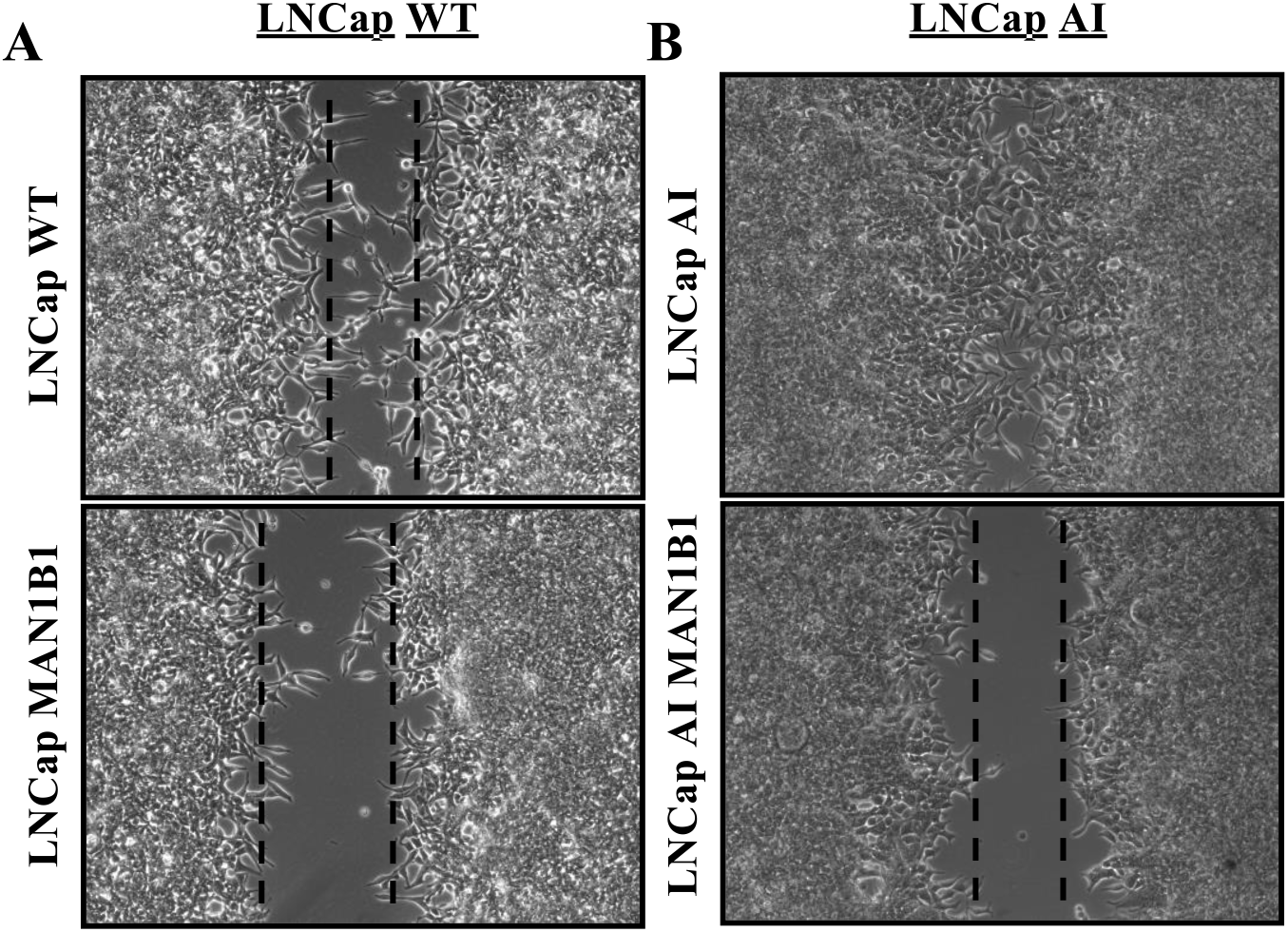
Overexpression of MAN1B1 in LNCaP (left) and LNCaP-AI (right) PCa cells alters the migratory phenotype. **A)** Scratch closure assay comparing LNCaP (WT) and LNCaP-MAN1B1 cell lines. LNCaP-MAN1B1 exhibited a modest reduction in the scratch closure rate compared to LNCaP (WT) cells. **B)** Scratch closure assay comparing LNCaP-AI (WT) and LNCaP-AI-MAN1B1 cell lines. LNCaP-MAN1B1 exhibited a reduction in the scratch closure rate compared to LNCaP-AI (WT) cells. Zone clearance for each panel is 500 μm.

### Downregulation of MAN1B1 expression in LNCaP PCa cells promotes an aggressive phenotype

Following the evaluation of the impact of MAN1B1 overexpression on the migratory phenotype of LNCaP, LNCaP-AI, and PC3 cell lines. We sought to assess the influence of MAN1B1 knock down (KD), to determine if it can induce a more aggressive PCa phenotype. We selected the LNCaP cell line as it was the least aggressive of the three cell lines of interest. LNCaP with reduced MAN1B1 expression was generated via siRNA knockdown (MAN1B1-siRNA LNCaP). Based on Western blot analysis, the expression of MAN1B1 in the LNCaP cell line was reduced in the MAN1B1-siRNA LNCaP cell line (Figure 4A) indicating the success of the MAN1B1 knockdown. To evaluate the impact of decreased MAN1B1 expression in LNCaP cells on migration capacity, wound healing assays were performed on LNCaP and LNCaP-MAN1B1-siRNA cell lines, and the migration potential was measured by the rate of cell motility. The silencing of MAN1B1 expression in LNCaP cells by siRNA had a profound impact on the behavior of LNCaP cells as shown by a wound healing assay (Figure 4B). LNCaP (NS) demonstrated low cellular motility over a 72 hr period as indicated by the lack of gap closure in the wound healing assay. On the contrary, knockdown of MAN1B1 in the LNCaP-siRNA MAN1B1 cells increased cellular motility, resulting in a near-complete healing of the cells in the wound healing assay (Figure 4B), suggesting that the reduced expression of MAN1B1 impacts cell migration, indicating a more aggressive PCa phenotype. Indeed, transwell migration analysis (Figure 4C) indicated a nearly two-fold increase in cellular mobility with MAN1B1 knockdown, underscoring the importance of MAN1B1 in promoting cellular adhesion.

**Figure 4.**
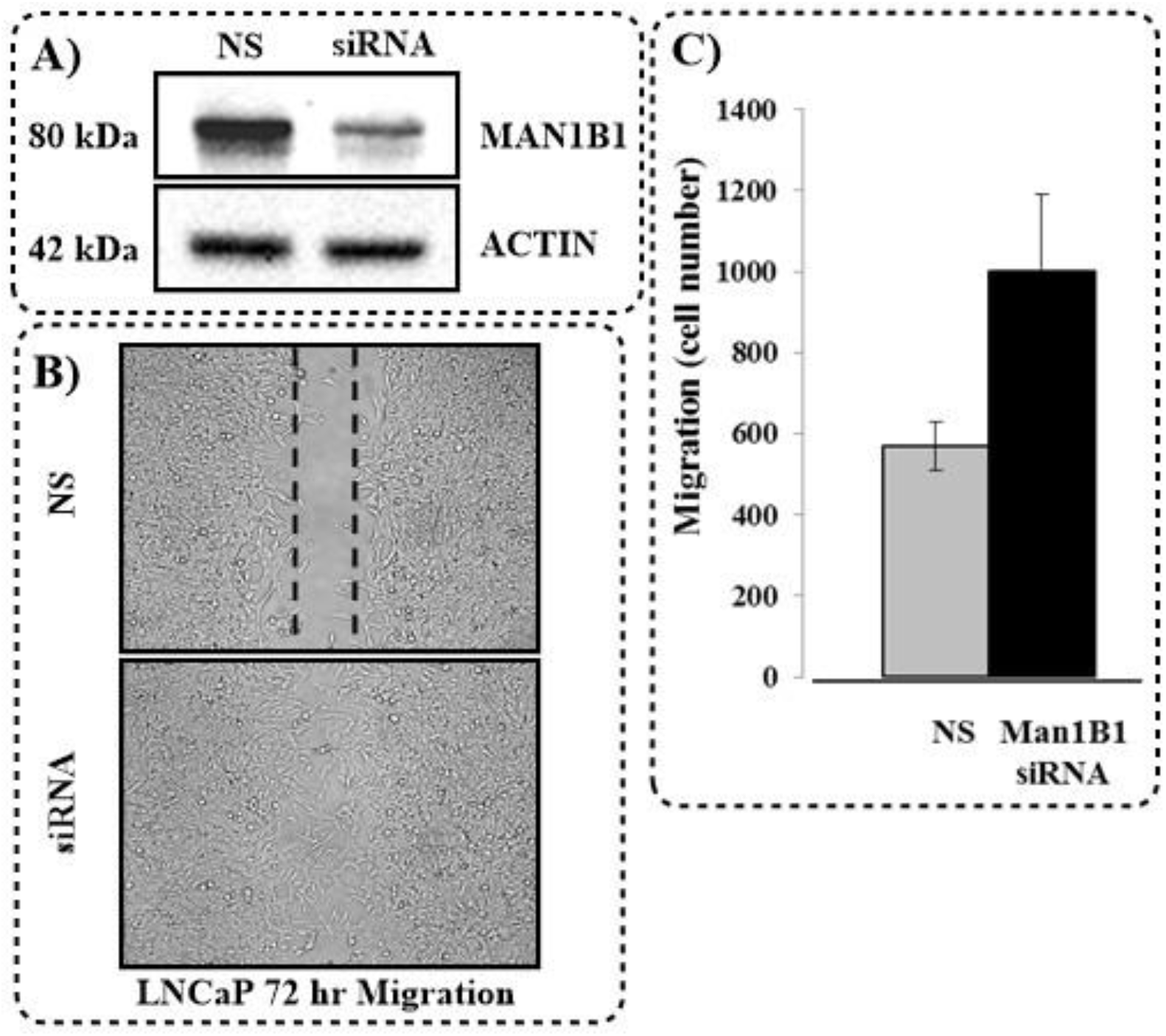
Downregulation of MAN1B1 expression in LNCaP cells drives LNCaP to a more aggressive phenotype. **A)** siRNA knockdown of MAN1B1 in LNCaP cells yielded a lower expression level compared to non-silenced control (NS). **B)** MAN1B1-siRNA LNCaP knockdown significantly enhances cell migration in wound healing. Zone clearance is 500 μm. **C)** Transwell migration after 24 hr indicates the enhanced migration efficiency of MAN1B1 knockdown LNCaP cells.

### Analysis of oligomannose glycans in LNCaP/LNCaP-MAN1B1 and LNCaP-AI/LNCaP-AI-MAN1B1 cell lines

In order to evaluate the impact of MAN1B1 on cellular glycosylation, we isolated N-glycans from the LNCaP, LNCaP-MAN1B1, LNCaP-AI, and LNCaP-AI-MAN1B1 cell lysates using the glycoprotein immobilization for glycan (GIG) extraction method previously reported by our group (34). The released glycans were purified using carbograph solid phase extraction (SPE) columns, and analyzed using matrix assisted laser desorption ionization with time of flight (MALDI-TOF). The relative peak intensity for each glycan was used as a relative estimation for glycan abundance, and the five major forms of high-mannose glycans were tabulated (Table 1). An increase in MAN1B1 expression would be expected to lead to a higher degree of glycan processing. We used the ratio between Man_8_GlcNAc_2_ and Man_9_GlcNAc_2_ (Man8/Man9) as an estimate of the degree of glycan processing, as a cell line with higher level of MAN1B1 expression should modify glycoproteins with Man_9_GlcNAc_2_ glycans to Man_8_GlcNAc_2_ in the ER (Figure 1). Indeed, in both LNCaP and LNCaP-AI cell lines, overexpression of MAN1B1 yielded higher ratios Man8/Man9 (Table 1). The glycomics analysis suggests that MAN1B1 overexpression results in higher glycan processing to Man_8_GlcNAc_2_ compared to WT cell lines.

**Table 1.** Glycomics analysis of LNCaP (WT), LNCaP (MAN1B1), LNCaP-AI (WT), and LNCaP-AI (MAN1B1) using MALDI-TOF analysis. The area under each peak was used to relatively quantify the fold-change values between WT cell lines and cell lines overexpressing MAN1B1.

| <u>Cell Line</u> | <u>Man5</u> | <u>Man6</u> | <u>Man7</u> | <u>Man8</u> | <u>Man9</u> | <u>Man8/Man9</u> |
| --- | --- | --- | --- | --- | --- | --- |
| LNCaP (WT) | 18955 | 31861 | 12234 | 12583 | 7221 | **1.7** |
| LNCaP (MAN1B1) | 10840 | 19461 | 7997 | 11315 | 1397 | **8.1** |
| LNCaP AI (WT) | 5086 | 13531 | 6561 | 9916 | 6197 | **1.6** |
| LNCaP AI (MAN1B1) | 3853 | 12177 | 6336 | 12606 | 2369 | **5.3** |

### Quantitative analysis of proteomic changes in MAN1B1 overexpressed PCa cells reveals downregulation of vimentin

The observation that MAN1B1 overexpression reduces cellular migration in three PCa cell lines, suggests that MAN1B1 overexpression can increase cellular adhesion, minimizing migratory capacity. Alternatively, MAN1B1 may have a pleiotropic regulatory role in EMT related proteins. To evaluate the pathways impacted by MAN1B1, leading to regulation in cellular migration and proliferation, changes in proteomic profiles were globally analyzed. In order to investigate the impact of MAN1B1 overexpression on LNCaP sublines, we performed proteomic analysis using 4-plex iTRAQ labelling, indicated changes in protein expression levels between PCa cell lines with and without MAN1B1 overexpression.

Several EMT related proteins including E-cadherin and vimentin were found by global proteomic analysis. E-cadherin (CDH1) is an important cell adhesion molecule that is normally expressed by epithelial cells. The abrogation or downregulation of E-cadherin expression is associated with an increase in the metastatic potential and invasiveness of numerous cancer cells, including PCa cells (35–39). E-cadherin is one of the hallmark EMT proteins that indicates a change in the migration phenotype during tumorigenesis. Given the central role of E-cadherin downregulation in facilitating downstream tumorigenesis, we analyzed the impact of MAN1B1 overexpression on the expression levels of E-cadherin. Quantitative proteomics indicated no change in E-cadherin expression in either of the three cell lines following MAN1B1 overexpression (Table 2). Similarly, no significant change in E-cadherin expression levels were observed in either LNCaP or LNCaPAI upon MAN1B1 overexpression (Table 2).

**Table 2.**
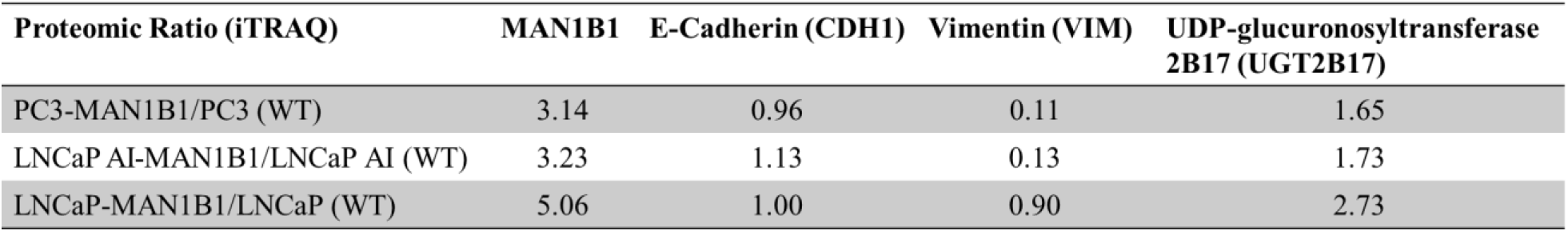
Proteomic results for protein expression of select proteins of interest in three PCa cell lines. Quantitative iTRAQ ratios are based on the iTRAQ channel intensity of MAN1B1 OE versus WT.

Conversely, Proteomic data demonstrated a dramatic drop in vimentin expression in both androgen-dependent PCa cell lines, LNCaP AI and PC3, with MAN1B1 overexpression, with no difference detected for androgen-independent PCa cell line, LNCaP (Table 2). The data suggests that overexpression of MAN1B1 in androgen-dependent PCa cell lines is sufficient to downregulate vimentin expression in LNCaP AI and PC3 cells, and promote a less aggressive phenotype, as overexpression of MAN1B1 is inversely correlated with vimentin expression (Table 2).

### Overexpression of MAN1B1 increases catabolism of androgen receptor (AR) agonists via increased UDP-glucuronosyltransferase 2B17 expression

In addition to the role of MAN1B1 in suppressing a critical EMT protein, vimentin, expression, a number of proximal tumor-repressor functions were observed. Another of the most striking observations was the concomitant upregulation in uridine diphosphate (UDP)-glucuronosyltransferase 2B17 (UGT2B17) expression with MAN1B1 overexpression. UGT2B17 an enzyme responsible for the glucuronosylation and subsequent clearance of potentially toxic xenobiotic and endogenous hydrophobic compounds (40). UGT2B17 expression in prostate tissue functions in the clearance of cellular androgens, and has been implicated in the progression of castration resistant PCa (CRPC) via promotion of ligand independent AR signaling (41). In quantitative proteomic analysis, MAN1B1 overexpression in LNCaP leads to a 2.7-fold increase in protein expression (Table 2). A similar, more modest phenotype was observed for LNCaP AI and LNCaP AI-MAN1B1, with a 1.73-fold change in expression based on proteomic data (Table 2). Similarly, quantitative proteomic analysis indicated a 1.65-fold increase in UGT2B17 expression in PC3-MAN1B1 cells compared to PC3 cells (Table 2).

## Discussion

The crucial steps in the EMT progression of tumor cells from a minimally aggressive to an aggressive phenotype are critically dependent upon the deregulation of both the cytoskeleton structure and the adhesion machinery. Both deregulation steps are required to allow detachment of the individual cells from the bulk tumor to seed their invasion to both the surrounding tissue and distant exogenous tissue. The overexpression of MAN1B1 in PCa cell lines reported herein has several functional consequences. First, the MAN1B1 expression appears to have an inverse correlation with PCa aggressiveness, such that downregulation of MAN1B1 expression induces a more aggressive PCa cell phenotype. Second, the downregulation of MAN1B1 leads to a number of pleiotropic and highly functionally significant alterations in the PCa cell proteome, most of which are amenable to promoting increased metastatic potential, decreased cellular senescence, and increased cellular proliferation.

In the present study, we have demonstrate that overexpression of MAN1B1 alone in LNCaP, LNCaP AI, and PC3 cells is sufficient to initiate the development of a non-aggressive phenotype, as indicated by the reduced cellular migratory capacity. Not only did we observe that overexpression of MAN1B1 decreased migration across all three cell lines utilized in this study, but siRNA-mediated knockdown of MAN1B1 in LNCaP was sufficient to induce higher migration potential and a more aggressive phenotype. The mechanism whereby MAN1B1 decreases PCa cellular migration is pleiotropic with a number of proteins related to the formation of adhesion junctions dysregulated, coinciding with MAN1B1 downregulation. Among these, expression of vimentin was found to be inversely correlated with MAN1B1 expression in both LNCaP AI and PC3 cells. It is interesting to note that none of the reported regulators of E-cadherin expression, Snail (42, 43), Slug (44), ZEB-1/2 (45), E12/E47 (46), or Twist (47) were found to be significantly altered in any of the cell lines that were utilized, suggesting a MAN1B1 regulation pathway that is independent of the known transcriptional regulators. Beyond the role of vimentin downregulation, the exact mechanism of MAN1B1 regulation of the EMT phenotype is less clear. The possibility that MAN1B1 regulates cell motility via post-transcriptional changes cannot be overstated. All of the major EMT-related proteins that were found to be altered in this study are known glycoproteins and it is possible that MAN1B1 modulates cell-cell adhesion through a partially transcriptionally-independent pathway, based primarily on the changes in protein glycosylation. The changes in the glycosylation profile of the cells upon MAN1B1 overexpression that were observed in this study could have a significant downstream impact on cell adhesion, such that the increased display of oligomannose glycans on MAN1B1-downregulated PCa cells promotes migration and invasiveness. Additional studies will be required to determine the specific contribution of post-translational modifications versus transcriptional modifications resulting from MAN1B1 overexpression. The complete absence of MAN1B1 expression results in a known type II congenital disorder of glycosylation (CDG) resulting in a devastating neurological phenotype characterized by significantly reduced cognitive capacity, obesity, dysmorphia, and is known to result in changes in Golgi morphology (48–50), underscoring the critical role of MAN1B1 in facilitating normal cellular function.

In conclusion, we have shown that the glycogene MAN1B1, is a potent pleiotropic regulator of PCa cell motility, where overexpression of MAN1B1 might suppress the invasive property of prostate cancer cells. Upregulation of MAN1B1 expression reduces LNCaP, LNCaP AI, and PC3 cell migration and restored a less-aggressive phenotype, consistent with an inhibitory function for MAN1B1. The dramatic phenotypic switch that was observed by increasing MAN1B1 expression suggests that it may be an important target of therapeutic design. As a non-toxic gene, upregulation of MAN1B1 expression may be a viable gene therapy target to slow the progression of advanced, metastatic PCa.

## Materials and Methods

### General

Unless otherwise specified, all chemicals and reagents were purchased from SigmaAldrich (St. Louis, MO, USA). All HPLC and LC-MS/MS solvents were obtained from Fisher Scientific (Pittsburgh, PA, USA).

### Cell culturing conditions

Human prostate cancer cell lines PC3 and LNCaP were purchased from ATCC (Manassas, VA). The LNCaP AI cell line was a gift from Dr. Steven Bova. The PCa cells were maintained in F-12K media with 10% HI fetal bovine serum (FBS) (Gibco, Grand Island, NY). Once reaching 80-90% cellular confluence, the cells were washed six times with ice-cold phosphate buffered saline, lysed and used from proteomic analysis.

### Establishment of stable expression of MAN1B1 in LNCaP, LNCaP AI, and PC3 cell lines

The Myc-DDK-tagged ORF clone of Homo Sapiens MAN1B1 or pCMV6-Entry (RC202824 and PS100001, respectively, Origene, Rockville, MD) vectors were transfected in LNCaP, LNCaP AI, or PC3 cell lines using Lipofectamine 2000 (Invitrogen) according to the manufacturer’s specifications. G418 (Invitrogen) at 400 μg/mL was used to screen the generated positive clones. The cell culture media was exchanged every three days during a period of two weeks. Once colonies formed, the cells from each clone was expanded for use in Western blot, genomic, proteomic, or migration analysis.

### MAN1B1 knockdown in LNCaP cells via siRNA

MAN1B1 knockdown was achieved using MAN1B1-specific short interfering RNA. In a 6-well plate, 12 μL of 20 μmol/L negative control siRNA (QIAGEN, Germantown, MD) or Hs_MAN1B1 siRNA (QIAGEN, Germantown, MD) was added to 88 μL of Opti-MEM reduced-serum medium (Thermo Fisher). Oligofectamine (12 μL) in 88 μL Opti-MEM medium was added to each well and the samples were incubated at RT for 25 min. The siRNA solution was added to cultured cells at 40-50% confluence in 800 μL of Opti-MEM media, and the cells were transfected for three hrs. Regular media containing 20% FBS, 1 mL total, was added to the transfected cells and the cells were further incubated for two day, before harvesting. Harvested cells were used for subsequent Western blot, genomic, and proteomic, and cell migration analysis.

### Western blot analysis

Western blotting was performed according to an established procedure (51). Briefly, PCa cells were lysed in 1X RIPA buffer (Millipore) and centrifuged at 15,000 x g for 15 min. Protein concentration was estimated using a Bradford colorimetric assay (BCA) kit (Thermo, Rockville, IL). Equal quantities (20 μg) of protein were loaded onto a 4-12% NuPAGE gel (Invitrogen, Carlsbad, CA). After running the gel, the proteins were applied to nitrocellulose membrane (Invitrogen). In order to positively identify the proteins, the following antibodies were used: MAN1B1 (SC-100543, 1:500) and actin (1:2000) (Cell Signaling Technology, Beverly, MA). Horse radish peroxidase (HRP)-conjugated secondary antibodies were incubated with the membrane at RT for 1 hr. Peroxidase activity was detected using X-ray film with an enhanced chemiluminescence detection system.

### Wound healing assay

Gap closure assays were performed in either an Ibidi Culture-Insert “wound healing” system, or in 6 mm Biocoat cell culture control inserts (BD Biosciences, San Jose, CA, USA). LNCaP, LNCaP siRNA-MAN1B1, LNCaP-MAN1B1, LNCaP AI, LNCaP AI-MAN1B1, PC3, or PC3-MAN1B1 cells (2 × 10^5^ average cells in 200 μL 0.2% FBS media) were seeded in the well containing 10% FBS until the cells reached confluence. An approximately 500 μm wide gap was generated by scratching the bottom of the well with a 200 μL pipette tip. The cells were incubated for 72 hrs at 37°C. Phase-contrast images of the wound healing process was obtained using a microscope with a 10× objective lens at 0 and 72 hrs after the scratch was made. The gap closure was compared between the initial 0 hr gap. Migrated cells were stained using Diff-Quik stain (Fisher Scientific, Pittsburgh, PA, USA).

### Transwell Assay

Transwell assay was performed using a 6 mm Biocoat cell culture control inserts (BD Biosciences, San Jose, CA). Cells were starved (overnight) in 0.2% FBS and PC3 cells (2 × 10^5^) were seeded into the upper well of a transwell chamber. To the bottom well of the chamber, 0.6 mL of 10% FBS was added. The cells were incubated for 24 hrs at 37°C. Migrated cells were stained with the Diff-Quik stain kit (Fisher Scientific, Pittsburg, PA). All cells were counted under microscope.

### Proliferation assay of PC3 cell lines

Approximately 1 × 10^4^ cells for both PC3 and PC3-MAN1B1 were seeded into a 96-well microtiter plate in 100 μL of culture medium with 10% FBS for 12 hrs or 144 hrs at 37°C in a 5% CO_2_ atmosphere. Following incubation, the number of living cells was subsequently measured using a 3-(4, 5-dimethylthiazol-2-yl)-5-(3-carboxymethoxyphenyl)-2-(4-sulfophenyl)-2H-tetrazolium (MTS) assay (Celltiter 96 Aqueous One-solution Cell Proliferation Assay, Promega, Madison, WI, USA) according to the manufacturer’s instructions. Absorbance of the wells at 490 nm was measured using a multiwell plate reader (Epoch, BioTek, Winooski, VT). Cell-free wells containing media alone served as blank controls. Each experiment was completed in eight replicates.

### Proteomic analysis using iTRAQ labeling

Cell pellets from LNCaP, LNCaP-MAN1B1, LNCaP AI, LNCaP AI-MAN1B1, PC3, and PC3-MAN1B1 were simultaneously lysed and denatured via sonication in 1 mL of 8 M urea containing 0.4 M ammonium bicarbonate (pH 8.0). Protein concentration was estimated by BCA protein assay (Thermo Fisher Scientific). Free cysteines were reduced in 120 mM tris(2-carboxyethyl)phosphine (TCEP) and alkylated by the addition of 160 mM iodoacetamide (IAA) at RT for 30 min. in the dark. The reduced and alkylated sample was diluted in 100 mM Tris-HCl, pH 7.5, containing 0.5 μg/μL trypsin. The digest was incubated at 37°C overnight. Digestion efficiency was measured via SDS-PAGE analysis, followed by silver staining. Digested peptides were purified with C18 desalting columns. Eluted peptides were dried using a speedvac.

To each cell line, containing 1 mg digested peptides, was added 250 μL iTRAQ dissolution buffer. Each iTRAQ (isobaric tags for relative and absolute quantitation) 4-plex channel was dissolved in 70 μL methanol and mixed with the peptide solution and incubated for 1 hr at RT. Two iTRAQ 4-plex sets were prepared. In both sets, channel 114 was used to label peptides from LNCaP WT cells. In the AI LNCaP iTRAQ analysis (set 1), channel 116 was used to label peptides from LNCaP AI WT cells, while channel 117 was used to label peptides from LNCaP AI-MAN1B1 cells. Similarly, in the PC3 iTRAQ analysis (set 2), channel 116 was assigned to PC3, while channel 117 was assigned to PC3-MAN1B1. Following iTRAQ labelling, the channels were pooled, desalted (C18 solid-phase extraction [SPE]), and purified by strong-cation exchange (SCX) SPE. 5% (approximately 100 μg) of the labelled peptides were dried and resuspended in 0.4% acetic acid prior to basic reverse-phase separation.

### Basic reverse phase liquid chromatography (bRPLC) fractionation

iTRAQ-labeled peptides were separated via bRPLC fractionation using a 1220 Infinity LC system, equipped with a Zorbax extended C18 analytical column (1.8 μm particle size, 4.6 × 100 mm; Agilent Technologies, Inc. CA, USA). The flow rate was set to 0.2 mL/min and a linear gradient of 92% A/8% B to 65% A/35% B in 85 min. was used to elute the peptide fractions. Buffer A consisted of 10 mM ammonium formate, pH 10, while buffer B was composed of 10 mM ammonium formate in 90% acetonitrile, pH 10. The eluted peptides were collected, concatenated into 24 fractions, and dried by speedvac.

### LC-MS/MS analysis

LC-MS/MS analysis of iTRAQ-labeled tryptic peptide fractions was performed using an LTQ-Orbitrap Velos (Thermo Fisher Scientific, Waltham, MA). Tryptic peptides were separated prior to analysis using a Dionex Ultimate 3000 RSL C nanosystem (Thermo Fisher Scientific) with a C18 PepMap RSLC column (75 μm × 50 cm). The analytical column was protected by a 5 mm guard column (Thermo Fisher Scientific). Chromatographic separation was performed at 250 nL/min during a linear gradient beginning from 95% A/5% B to 68% A/32% B in 90 min. Solvent A consisted of 0.1% formic acid (FA) in water, and B consisted of 90% aqueous ACN containing 0.1% FA. The Orbitrap Velos was operating in high-energy collision-induced dissociation (CID) mode with the normalized collision energy set to 45.

### Oligomannose analyses of prostate cancer cells with Man1B1 overexpression

Glycomics analysis was performed using the glycoprotein immobilization for glycan extraction (GIG) method reported previously (34, 52, 53). N-linked glycans released from glycoproteins using Peptide:N-glycosidase F (PNGase F, New England Biolabs, Ipswich, MA) were mixed with 2,5-dihydroxybenzoic acid (DHB) containing dimethylaniline and spotted on a 384-well μFocus matrix assisted laser desorption/ionization (MALDI) plate (Hudson Surface Technology, Fort Lee, NJ). Released glycans were analyzed using a Shimadzu AXIMA Resonance MALDI quadrupole ion trap-time-of-flight (MALDI-QIT-TOF) mass spectrometer (Shimadzu, Columbia, MD) in positive ionization mode.

### MS data analysis

The acquired MS/MS spectra was searched directly using the SEQUEST search engine within Proteome Discoverer (PD) 1.4.0.288 (Thermo Fisher Scientific). The database used was the NCBI homo sapien FASTA file database, using a 1% protein false discovery rate (FDR). The lone fixed modification was carbamidomethylation of reduced cysteine residues, while oxidation of methionine was set as a variable modification. iTRAQ 4-plex labeling of the N-termini and lysine residues was set as dynamic modifications. The search tolerated a maximum of two missed cleavages for trypsin digestion. The precursor mass tolerance was 10 ppm, while fragment ion mass tolerance was set to 20 ppm. Quantitation of the iTRAQ intensity signal was performed using peptide intensity in PD, and all median peptides intensities were normalized in PD. The iTRAQ ratio cutoff for upregulation/downregulation was 2-fold and 0.5-fold, respectively.

## Acknowledgements

This research was supported by the Early Detection Research Network (EDRN), an initiative of the National Cancer Institute (NCI U01CA152813).

The mass spectrometry proteomics data have been deposited to the ProteomeXchange Consortium via the PRIDE (54) partner repository with the dataset identifier PXD015201.

## References

1. Siegel, R., Ma, J., Zou, Z., and Jemal, A. (2014) Cancer statistics, 2014 - Siegel - 2014 - CA: A Cancer Journal for Clinicians - Wiley Online Library. … a Cancer J. Clin. Volume 65, Issue 1 January/February 2015 Pages 5–29

2. Rider, J. R., Sandin, F., Andrén, O., Wiklund, P., Hugosson, J., and Stattin, P. (2013) Long-term outcomes among noncuratively treated men according to prostate cancer risk category in a nationwide, population-based study. Eur. Urol. 63, 88–96

3. Pokala, N., Huynh, D. L., Henderson, A. A., and Johans, C. (2015) Survival Outcomes in Men Undergoing Radical Prostatectomy After Primary Radiation Treatment for Adenocarcinoma of the Prostate. Clin. Genitourin. Cancer. 10.1016/j.clgc.2015.12.010

4. Heinlein, C. A., and Chang, C. (2004) Androgen receptor in prostate cancer. Endocr. Rev. 25, 276–308

5. Chen, Y., Sawyers, C. L., and Scher, H. I. (2008) Targeting the androgen receptor pathway in prostate cancer. Curr. Opin. Pharmacol. 8, 440–448

6. Roca, H., Jones, J. D., Purica, M. C., Weidner, S., Koh, A. J., Kuo, R., Wilkinson, J. E., Wang, Y., Daignault-Newton, S., Pienta, K. J., Morgan, T. M., Keller, E. T., Nör, J. E., Shea, L. D., and McCauley, L. K. (2017) Apoptosis-induced CXCL5 accelerates inflammation and growth of prostate tumor metastases in bone. J. Clin. Invest. 10.1172/JCI92466

7. Goel, H. L., Li, J., Kogan, S., and Languino, L. R. (2008) Integrins in prostate cancer progression. Endocr. Relat. Cancer. 15, 657–664

8. Kuo, P. L., Chen, Y. H., Chen, T. C., Shen, K. H., and Hsu, Y. L. (2011) CXCL5/ENA78 increased cell migration and epithelial-to-mesenchymal transition of hormone-independent prostate cancer by early growth response-1/snail signaling pathway. J. Cell. Physiol. 226, 1224–1231

9. Liotta, L. A., Steeg, P. S., and Stetler-Stevenson, W. G. (1991) Cancer metastasis and angiogenesis: An imbalance of positive and negative regulation. Cell. 64, 327–336

10. Guo, W., and Giancotti, F. G. (2004) Integrin signalling during tumour progression. Nat. Rev. Mol. Cell Biol. 5, 816–826

11. M, D. M., and Baluk, P. (2002) Significance of blood vessel leakiness in cancer. Cancer Res. 62, 5381–5385

12. Hashizume, H., Baluk, P., Morikawa, S., McLean, J. W., Thurston, G., Roberge, S., Jain, R. K., and McDonald, D. M. (2000) Openings between defective endothelial cells explain tumor vessel leakiness. Am. J. Pathol. 156, 1363–1380

13. Liu, C.-Y., Lin, H.-H., Tang, M.-J., and Wang, Y.-K. (2015) Vimentin contributes to epithelial-mesenchymal transition cancer cell mechanics by mediating cytoskeletal organization and focal adhesion maturation. Oncotarget. 6, 15966–15983

14. Stowell, S. R., Ju, T., and Cummings, R. D. (2015) Protein Glycosylation in Cancer. Annu. Rev. Pathol. Mech. Dis. 10, 473–510

15. Meany, D. L., and Chan, D. W. (2011) Aberrant glycosylation associated with enzymes as cancer biomarkers. Clin. Proteomics. 10.1186/1559-0275-8-7

16. Padler-Karavani, V. (2014) Aiming at the sweet side of cancer: Aberrant glycosylation as possible target for personalized-medicine. Cancer Lett. 352, 102–112

17. Gilgunn, S., Conroy, P. J., Saldova, R., Rudd, P. M., and O’Kennedy, R. J. (2013) Aberrant PSA glycosylation - A sweet predictor of prostate cancer. Nat. Rev. Urol. 10, 99–107

18. Chen, C.-Y., Jan, Y.-H., Juan, Y.-H., Yang, C.-J., Huang, M.-S., Yu, C.-J., Yang, P.-C., Hsiao, M., Hsu, T.-L., and Wong, C.-H. (2013) Fucosyltransferase 8 as a functional regulator of nonsmall cell lung cancer. Proc. Natl. Acad. Sci. 110, 630–635

19. Wang, X., Chen, J., Li, Q. K., Peskoe, S. B., Zhang, B., Choi, C., Platz, E. A., and Zhang, H. (2014) Overexpression of α (1,6) fucosyltransferase associated with aggressive prostate cancer. Glycobiology. 24, 935–944

20. de Leoz, M. L. A., Young, L. J. T., An, H. J., Kronewitter, S. R., Kim, J., Miyamoto, S., Borowsky, A. D., Chew, H. K., and Lebrilla, C. B. (2011) High-Mannose Glycans are Elevated during Breast Cancer Progression. Mol. Cell. Proteomics. 10, M110.002717

21. Maverakis, E., Kim, K., Shimoda, M., Gershwin, M. E., Patel, F., Wilken, R., Raychaudhuri, S., Ruhaak, L. R., and Lebrilla, C. B. (2015) Glycans in the immune system and The Altered Glycan Theory of Autoimmunity: A critical review. J. Autoimmun. 57, 1–13

22. Talabnin, K., Talabnin, C., Ishihara, M., & Azadi, P. (2018) Increased expression of the high‑mannose M6N2 and NeuAc3H3N3M3N2F tri‑antennary N‑glycans in cholangiocarcinoma. Oncol. Lett. 15, 1030–1036

23. Chen, H., Deng, Z., Huang, C., Wu, H., Zhao, X., and Li, Y. (2017) Mass spectrometric profiling reveals association of *N*-glycan patterns with epithelial ovarian cancer progression. Tumor Biol. 39, 101042831771624

24. Balog, C. I. a, Stavenhagen, K., Fung, W. L. J., Koeleman, C. a, McDonnell, L. a, Verhoeven, A., Mesker, W. E., Tollenaar, R. a E. M., Deelder, A. M., and Wuhrer, M. (2012) N-glycosylation of colorectal cancer tissues: a liquid chromatography and mass spectrometry-based investigation. Mol. Cell. Proteomics. 11, 571–85

25. Sethi, M. K., Thaysen-Andersen, M., Smith, J. T., Baker, M. S., Packer, N. H., Hancock, S., and Fanayan, S. (2014) Comparative N-glycan profiling of colorectal cancer cell lines reveals unique bisecting GlcNAc and ??-2,3-linked sialic acid determinants are associated with membrane proteins of the more metastatic/aggressive cell lines. J. Proteome Res. 13, 277–288

26. Tabarés, G., Radcliffe, C. M., Barrabés, S., Ramírez, M., Aleixandre, N., Hoesel, W., Dwek, R. A., Rudd, P. M., Peracaula, R., and de Llorens, R. (2006) Different glycan structures in prostate-specific antigen from prostate cancer sera in relation to seminal plasma PSA. Glycobiology. 16, 132–145

27. Wang, D., Dafik, L., Nolley, R., Huang, W., Wolfinger, R. D., Wang, L. X., and Peehl, D. M. (2013) Anti-oligomannose antibodies as potential serum biomarkers of aggressive prostate cancer. Drug Dev. Res. 74, 65–80

28. Kornfeld, R. (1985) Assembly of Asparagine-Linked Oligosaccharides. Annu. Rev. Biochem. 54, 631–664

29. Jin, Z.-C., Kitajima, T., Dong, W., Huang, Y.-F., Ren, W.-W., Guan, F., Chiba, Y., Gao, X.-D., and Fujita, M. (2018) Genetic disruption of multiple α1,2-mannosidases generates mammalian cells producing recombinant proteins with high-mannose–type N-glycans. J. Biol. Chem. . 293, 5572–5584

30. Phoomak, C., Silsirivanit, A., Park, D., Sawanyawisuth, K., Vaeteewoottacharn, K., Wongkham, C., Lam, E. W.-F., Pairojkul, C., Lebrilla, C. B., and Wongkham, S. (2018) O-GlcNAcylation mediates metastasis of cholangiocarcinoma through FOXO3 and MAN1A1. Oncogene. 10.1038/s41388-018-0366-1

31. Pulukuri, S. M. K., Gondi, C. S., Lakka, S. S., Jutla, A., Estes, N., Gujrati, M., and Rao, J. S. (2005) RNA interference-directed knockdown of urokinase plasminogen activator and urokinase plasminogen activator receptor inhibits prostate cancer cell invasion, survival, and tumorigenicity in vivo. J. Biol. Chem. 280, 36529–36540

32. Dalrymple, S., Antony, L., Xu, Y., Uzgare, A. R., Arnold, J. T., Savaugeot, J., Sokoll, L. J., De Marzo, A. M., and Isaacs, J. T. (2005) Role of Notch-1 and E-cadherin in the differential response to calcium in culturing normal versus malignant prostate cells. Cancer Res. 65, 9269–9279

33. Yu, P., Duan, X., Cheng, Y., Liu, C., Chen, Y., Liu, W., Yin, B., Wang, X., and Tao, Z. (2017) Androgen-independent LNCaP cells are a subline of LNCaP cells with a more aggressive phenotype and androgen suppresses their growth by inducing cell cycle arrest at the G1 phase. Int. J. Mol. Med. 40, 1426–1434

34. Shah, P., Yang, S., Sun, S., Aiyetan, P., Yarema, K. J., and Zhang, H. (2013) Mass spectrometric analysis of sialylated glycans with use of solid-phase labeling of sialic acids. Anal. Chem. 10.1021/ac3033867

35. Davies, G., Jiang, W. G., and Mason, M. D. (2000) Cell-cell adhesion molecules and signaling intermediates and their role in the invasive potential of prostate cancer cells. J. Urol. 163, 985–992

36. Annicotte, J. S., Iankova, I., Miard, S., Fritz, V., Sarruf, D., Abella, A., Berthe, M. L., Noel, D., Pillon, A., Iborra, F., Dubus, P., Maudelonde, T., Culine, S., and Fajas, L. (2006) Peroxisome proliferator-activated receptor {gamma} regulates E-cadherin expression and inhibits growth and invasion of prostate cancer. Mol.Cell Biol. 26, 7561–7574

37. Bae, K.-M., Parker, N. N., Dai, Y., Vieweg, J., and Siemann, D. W. (2011) E-cadherin plasticity in prostate cancer stem cell invasion. Am. J. Cancer Res. 1, 71–84

38. Umbas, R., Isaacs, W. B., Bringuier, P. P., Schaafsma, H. E., Karthaus, H. F., Oosterhof, G. O., Debruyne, F. M., and Schalken, J. A. (1994) Decreased E-cadherin expression is associated with poor prognosis in patients with prostate cancer. Cancer Res. 54, 3929–3933

39. Umbas, R., Isaacs, W. B., Bringuier, P. P., Xue, Y., Debrune, F. M. J., and Schalken, J. A. (1997) Relation between aberrant ??-catenin expression and loss of E-cadherin function in prostate cancer. Int. J. Cancer. 74, 374–377

40. Park, J. Y., Tanner, J. P., Sellers, T. A., Huang, Y., Stevens, C. K., Dossett, N., Shankar, R. A., Zachariah, B., Heysek, R., and Pow-Sang, J. (2007) Association Between Polymorphisms in HSD3B1 and UGT2B17 and Prostate Cancer Risk. Urology. 70, 374–379

41. Li, H., Xie, N., Chen, R., Verreault, M., Fazli, L., Gleave, M. E., Barbier, O., and Dong, (2016) UGT2B17 expedites progression of castration-resistant prostate cancers by promoting ligand-independent AR signaling. Cancer Res. 76, 6701–6711

42. Batlle, E., Sancho, E., Francí, C., Domínguez, D., Monfar, M., Baulida, J., and De Herreros, A. G. (2000) The transcription factor Snail is a repressor of E-cadherin gene expression in epithelial tumour cells. Nat. Cell Biol. 2, 84–89

43. Cano, A., Pérez-Moreno, M. A., Rodrigo, I., Locascio, A., Blanco, M. J., Del Barrio, M. G., Portillo, F., and Nieto, M. A. (2000) The transcription factor Snail controls epithelial-mesenchymal transitions by repressing E-cadherin expression. Nat. Cell Biol. 2, 76–83

44. Bolos, V., Peinado, H., Perez-Moreno, M. A., Fraga, M. F., Esteller, M., and Cano, A. (2016) The transcription factor Slug represses E-cadherin expression and induces epithelial to mesenchymal transitions: a comparison with Snail and E47 repressors. J. Cell Sci. 129, 1283–1283

45. Comijn, J., Berx, G., Vermassen, P., Verschueren, K., Van Grunsven, L., Bruyneel, E., Mareel, M., Huylebroeck, D., and Van Roy, F. (2001) The two-handed E box binding zinc finger protein SIP1 downregulates E-cadherin and induces invasion. Mol. Cell. 7, 1267–1278

46. Pérez-Moreno, M. A., Locascio, A., Rodrigo, I., Dhondt, G., Portillo, F., Nieto, M. A., and Cano, A. (2001) A New Role for E12/E47 in the Repression of E-cadherin Expression and Epithelial-Mesenchymal Transitions. J. Biol. Chem. 276, 27424–27431

47. Vesuna, F., van Diest, P., Chen, J. H., and Raman, V. (2008) Twist is a transcriptional repressor of E-cadherin gene expression in breast cancer. Biochem. Biophys. Res. Commun. 367, 235–241

48. Van Scherpenzeel, M., Timal, S., Rymen, D., Hoischen, A., Wuhrer, M., Hipgrave-Ederveen, A., Grunewald, S., Peanne, R., Saada, A., Edvardson, S., Grønborg, S., Ruijter, G., Kattentidt-Mouravieva, A., Brum, J. M., Freckmann, M. L., Tomkins, S., Jalan, A., Prochazkova, D., Ondruskova, N., Hansikova, H., Willemsen, M. A., Hensbergen, P. J., Matthijs, G., Wevers, R. A., Veltman, J. A., Morava, E., and Lefeber, D. J. (2014) Diagnostic serum glycosylation profile in patients with intellectual disability as a result of MAN1B1 deficiency. Brain. 137, 1030–1038

49. Rymen, D., Peanne, R., Millón, M. B., Race, V., Sturiale, L., Garozzo, D., Mills, P., Clayton, P., Asteggiano, C. G., Quelhas, D., Cansu, A., Martins, E., Nassogne, M. C., Gonçalves-Rocha, M., Topaloglu, H., Jaeken, J., Foulquier, F., and Matthijs, G. (2013) MAN1B1 Deficiency: An Unexpected CDG-II. PLoS Genet. 10.1371/journal.pgen.1003989

50. Rafiq, M. A., Kuss, A. W., Puettmann, L., Noor, A., Ramiah, A., Ali, G., Hu, H., Kerio, N. A., Xiang, Y., Garshasbi, M., Khan, M. A., Ishak, G. E., Weksberg, R., Ullmann, R., Tzschach, A., Kahrizi, K., Mahmood, K., Naeem, F., Ayub, M., Moremen, K. W., Vincent, J. B., Ropers, H. H., Ansar, M., and Najmabadi, H. (2011) Mutations in the alpha 1,2-mannosidase gene, MAN1B1, cause autosomal-recessive intellectual disability. Am. J. Hum. Genet. 89, 176–182

51. Wang, X., Gu, J., Ihara, H., Miyoshi, E., Honke, K., and Taniguchi, N. (2006) Core fucosylation regulates epidermal growth factor receptor-mediated intracellular signaling. J. Biol. Chem. 281, 2572–2577

52. Yang, S., Li, Y., Shah, P., and Zhang, H. (2013) Glycomic analysis using glycoprotein immobilization for glycan extraction. Anal. Chem. 85, 5555–5561

53. Yang, S., Höti, N., Yang, W., Liu, Y., Chen, L., Li, S., and Zhang, H. (2017) Simultaneous analyses of N-linked and O-linked glycans of ovarian cancer cells using solid-phase chemoenzymatic method. Clin. Proteomics. 10.1186/s12014-017-9137-1

54. Perez-Riverol, Y., Csordas, A., Bai, J., Bernal-Llinares, M., Hewapathirana, S., Kundu, D. J., Inuganti, A., Griss, J., Mayer, G., Eisenacher, M., Pérez, E., Uszkoreit, J., Pfeuffer, J., Sachsenberg, T., Yilmaz, Ş., Tiwary, S., Cox, J., Audain, E., Walzer, M., Jarnuczak, A. F., Ternent, T., Brazma, A., and Vizcaíno, J. A. (2019) The PRIDE database and related tools and resources in 2019: Improving support for quantification data. Nucleic Acids Res. 10.1093/nar/gky1106

